# Survival of environmental DNA in sediments: Mineralogic control on DNA taphonomy

**DOI:** 10.1101/2020.01.28.922997

**Authors:** C.L. Freeman, L. Dieudonné, O.B.A. Agbaje, M. Zure, J.Q. Sanz, M. Collins, K.K. Sand

## Abstract

Extraction of environmental DNA (eDNA) from sediments are providing ground-breaking views of the past ecosystems and biodiversity. Despite this rich source of information, it is still unclear which sediments favour preservation and why. Here we used atomic force microscopy and molecular dynamics simulations to explore the DNA-mineral interaction to assess how mineralogy and interfacial geochemistry play a role in the preservation of environmental DNA on mineral substrates. We demonstrate that mineral composition, surface topography and surface charge influence DNA adsorption behavior as well as preservation. Modeling and experimental data show that DNA damage can be induced by mineral binding if there is a strong driving force for adsorption. The study shows that knowledge of the mineralogical composition of a sediment and the environmental conditions can be useful for assessing if a deposit is capable of storing extracellular DNA and to what extent the DNA would be preserved. Our data adds to the understanding of eDNA taphonomy and highlights that, for some mineral systems, fragmented DNA may not represent old DNA.

## Introduction

Arguably the most significant breakthrough in the analysis of environmental sedimentary DNA is the ability to recover DNA sequences. ^1,2^ The perspectives of harvesting the environmental sedimentary genetic source are vast and include the possibility of recovering biodiversity across time and space ^3,4^ as well as identifying the presence of archaic humans ^5^ or record animal processing ^6^ in the absence of fossils. The range of sediments sampled targeted are quite diverse and include chemically precipitated sediments such as speleothems, ^7,8,9–11^ and depositional sediments such as marine, soil or lake sediments. The major component by mass in speleothems are carbonates and in the majority of the depositional sediments are silicates. Currently we try to identify sedimentary sites with preserved DNA based on the low potential for degradation by dynamic biological, physical, and chemical factors, ^12^ and exclude considerations of the processes and mechanism determining the DNA-mineral associations that drive retention and storage of DNA on mineral surface. In this study we use a bottom-up approach to explore DNA-mineral interactions with the focus on understanding the geochemical interplay between mineral surface composition and solution composition for DNA adsorption behaviour and stability. With this approach we introduce the mineralogy and environmental conditions as a tool for assessing the DNA preservation potential of a particular site.

Sediments display a wide variation in mineralogical composition which, considering interfacial geochemical principles, will influence their DNA adsorption capacity. Additionally, the environmental conditions, such as solution composition, is expected to play a role for both the adsorption process and for the subsequent preservation of the DNA. Essentially, to extract and analyse sedimentary DNA, we are reliant on the stabilisation of DNA by the sediment under specific environmental conditions. Not much, however, is known about how DNA is stabilized by sediments, which prevents us from identifying promising sites where DNA is stabilized. Minerals differ not only in composition and structure but also in charge density which affects the bond strength, the adsorption behavior as well as DNA adsorption capacity. The surface charge of most mineral surfaces changes as a function of pH where in general, silicates are negatively charged and oxides, carbonates and hydroxides are positively charged over a wide pH range. ^13^ DNA have been shown to mainly interact through its phosphate backbone ^14,15^ which is negatively charged above pH 5. The phosphate backbone can, through electrostatic forces, interact directly with positively charged surfaces, whereas the availability of polyvalent cations have been shown to determine its binding to a negatively charged surface. ^16,17,18–22^ The charge density of most mineral surfaces leads to the adsorption of tightly bound layers of water molecules ^23^ Geissbühler et al 2004, ^23^ and may promote hydrolysis of the DNA. ^24^ The DNA molecule can adsorb to the mineral via water layers or penetrate the water layers and interact with the mineral directly which would be expected to form a stronger bond. Most studies of DNA adsorption to minerals are conducted in bulk and we lack information on adsorption dynamics, associated conformation as well as a mechanistic understanding of the interaction bonds.

Atomic force microscopy (AFM) imaging has provided nano level information on DNA binding and conformation as a function of solution compositions on mica surfaces. ^25–27^ Mica (muscovite) is negatively charged and can be considered as an analogue to the basal planes of clay mineral surfaces and can be considered as a reasonable model for studying DNA behaviours on silicates. It is well established that the DNA conformation on mica changes with cationic content and ionic strength, where a larger ionic potential (charge/density) favours adsorption. This is classical behaviour of outer sphere bonding ^28^ which was recently confirmed by. ^29^ Because of the need for a cationic bridge to mediate the DNA-mica interaction, the adsorption will be sensitive to solution compositions and the preservation of the DNA may well be jeopardized if the environmental conditions changes. Indeed, DNA does show reversible bonding on mica, where pH or cationic composition drive the reactions. ^27^ Polymer-calcite associations have been widely studied using molecular dynamics MD simulations and AFM and the combination of the two techniques provide insight into the adsorption behaviour and enable an assessment of the underlying mechanisms. MD simulations offer mechanistic level access to the processes associated with binding, the bonds involved in the binding and their stabilities. ^30,31^

Speleothems from caves consist predominantly of calcite (CaCO3) and in contrast to mica, calcite is overall positively charged in most environmental conditions. Calcite has a heterogeneous surface distribution of charges and topography. The (10.4) face is the most commonly expressed and energetically stable face of natural calcite. The (10.4) face comprises atomically flat terraces that intersect at step edges. Because of calcite’s *orthorhombic and rhombohedral structure*, there are 2 inequivalent step edges: obtuse and acute which can each be terminated by carbonate or calcium ions and hence display locally positive or negative charges toward the solution. MD simulations have already provided much information on how interfacial water can control the binding of molecules at the calcite surface. Generally, it can be hard for molecules to penetrate the tightly bound surface water layers. ^32,33–35^ The presence of steps can be crucial here as these break up the structure of the water providing more accessible sites for molecular binding. ^32,36,37–39^ The large degree of intrinsic structure, high negative charge and lack of flexible binding groups makes DNA binding to calcite very different to previous simulations of proteins, peptides and other small molecules. Given the potential of calcite as a DNA archive it is important to understand the binding process.

Here we adsorbed circular DNA molecules (plasmid DNA with 2686 base pairs) onto calcite in the presence of three different ions with distinct ionic potentials and used AFM to compare the resulting DNA binding and conformation with those reported for mica. We chose plasmid DNA to exclude interaction from the base pairs in an open strand and to be compatible with the modeling that considered a closed strand in its starting configuration. We generated etch pits on the calcite surface to increase the amount of step edges on the calcite crystals during adsorption to understand if there was a preference of DNA for adsorbing to terrace or edge sites, and acute or obtuse steps. We used MD simulations to support the observed binding preferences as seen with the AFM and to understand the reason for them.

We subsequently stressed the adsorbed DNA by slightly dissolving the calcite surface to investigate plasmid fragmentation in non-equilibrium conditions. Combined, our results highlight the importance of the interplay between solution conditions and mineral surface charge for adsorbing and stabilising DNA. Our study increases our understanding of DNA taphonomy in the environment and opens for a new approach for assessing preservation potential of environmental DNA in various sedimentary systems.

## Materials and Methods

### Minerals

Single crystals of optical quality Iceland spar calcite (purchased from Ward’s Natural Science,USA) . The mica were grade V1, had a diameter of 1 cm and were purchased from Ted Pella Inc.

### Chemicals

The salts (NaCl, MgCl_2_, NiCl_2_) were reagent grade and purchased from Sigma Aldrich and used without purification. The Poly-L-Lysine was purchased from Boster Biological Technology and used as received. The DNA was the double stranded plasmid: pUC19. It is composed of 2686 base pairs and is approximately 910 nm long. The plasmid was purchased from Integrated DNA Technologies IDT. For imaging and sample preparation we used a nuclease free Tris buffer at pH 7,5 purchased from. All solutions were prepared using ultrapure water from a MilliQ deionizing column (resistivity > 18 MΩ cm; Millipore Corporation).

### Substrate preparation

Calcite samples were cleaved in air, and the calcite dust was immediately removed with a firm stream of N_2_(g). The mica was cleaved with a piece of tape. The Poly-L-Lysine film was prepared on freshly cleaved mica by using a 10 μL droplet of Poly-L-Lysine at a concentration of 0,01 w/v in MilliQ water. After 30 s the surface was rinsed with plenty of MilliQ and blow dried with N_2_(g)

### Preparation of DNA on the minerals

A 10 μL droplet of plasmid DNA pUC19 at a concentration of 0,4 ng/μL and 10 mM electrolyte was placed on a freshly cleaved mica or calcite. The droplet was left for two minutes before rinsed with 400 μL MilliQ. The mineral was dried with a soft blow of N_2_(g) for 30 seconds.

### AFM imaging

For scanning in air and liquids, we used a Cypher and a Cypher VRS from Oxford Instruments. Both were running in tapping mode. For the imaging in air we used aluminium coated AC240TS silicon tips from Olympus with spring constants between 0.6 and 3.5 N/m and a resonance frequency approximating 70 kHz. For tapping in liquid, we used BL-AC10DS from Asylum Research, Oxford Instruments. We used a spring constant of 107 pN/nm and a resonance frequency of 1500 kHz. The spring constants were determined using the Sader method ^40^. For both instruments, the force between the tip and the sample was varied to minimize the force exerted by the tip on the surface. Scan angle, scan rate, and set-point were systematically varied. We measured multiple sites on every sample and the experiments were repeated several times. For the fragmentation experiments we prepared the sample as for the experiments performed in air and imaged the surface in air to confirm that we had intact plasmids. Subsequently, we introduced the calcite saturated buffer to the surface and began the approach process.

### Chemical Force Microscopy (CFM) - Surface chemical analysis

After the tips were checked under a microscope, the tips were cleaned with water for molecular biology, ethanol and finally under ultraviolet ozone cleaner for 20 minutes. Subsequently, the tips were functionalized overnight with a self-assembled monolayer (SAM) solution terminating in phosphate (PO_4_^3-^) end group (Sigma-Aldrich, 754269). Then, the tips were washed in ethanol (3×), and allowed to dry on glass slide in a petri dish at room temperature. Like AFM imaging, we used a Cypher and a Cypher VRS from Oxford Instruments and BL-AC10DS from Asylum Research from Oxford Instruments for tapping in liquid. For these measurements, we used OBL-10 silicon nitride tips from Bruker with spring constants between 0.03 and 0.009 N/m and a resonance frequency approximating 70 kHz. A constant trigger force at 80 pN was used and the spring constants of the levers were determined by thermal noise method (Hutter and Bechhoefer 1993; Butt and Jaschke 1995). We recorded force-distance cycles at a 3 μm scan size, 300 nm force distance, 1 Hz scan rate, and both approach and retract velocities are approximating 1 μm/s. For the CFM force map, force lines and points are constant at 14N, and scan angle is at zero degree. We performed the measurements in liquid at concentrations of 10 mM (NaCl and MgCl_2_) and 100 mM (NaCl and MgCl_2_), pH 7. For each solution parameter, the experiment was repeated several times and average was calculated. A single tip of each type was used for tapping mode on the surfaces of freshly cleaved mica and calcite to ensure that the spring constant and tip radius were constant. At least we carried out two independent measurements to check the uniformity of our analysis.

### MD simulations

All molecular dynamics simulations were performed using DL_POLY classic (Smith and Forester 1996). Simulations were carried out in the NVT ensemble with a temperature of 300 K and a thermostat relaxation time of 0.1 ps. The leapfrog verlet algorithm was used with a timestep of 0.5 fs. Coulombic interactions were modelled with the Ewald method (precision 1×10^-6^). Periodic boundary conditions were used for all the simulations.

The calcite forcefield was taken from ^41^ using SPC/E water. The DNA forcefield was taken from the AMBER forcefield. ^42^ Water-DNA interactions were calculated using the standard Lorentz-Berthelot mixing rules. The Ca-N interactions were taken from Freeman et al 2009, Ca-O (in DNA) were fitted using the Schröder method as described in ^43^. Terms are listed in supplementary material.

The DNA molecule was a random double helix with the sequence C(G)-A(T)-A(T)-G(C)-T(A)-T(A)- T(A)-A(A)-G(C)-C(G)-T(A)-, where A is adenine, T is Thymine, G is guanine and C is cytosine. The complementary base is listed in brackets. The periodic simulation cell was constructed such that the z cell parameter size matched the chain length of the 11 base pair chain (39.5 Å). This meant the DNA chain periodically spanned the boundary conditions (i.e. the end C(G) connected with the other end T(A)). The calcite slab was built in Metadise (Watson et al 1996) from a geometry optimised cell in GULP ^44^ The slab was cut to express the (10.4) surface of calcite. Steps were added by removing Ca and CO3 ions from one surface and placing them on the other surface in the appropriate lattice positions. After all surface cuts and reorganisations the calcite slab was relaxed in vacuum. The final cells consisted of 8 CaCO_3_ layers with 768 formula units with surface vectors 47.8 Å × 39.5 Å.

For the adsorption studies the DNA was placed parallel to the surface with the centre of mass ~11 Å from the calcite surface. 1268 water molecules and 11 Ca^2+^ cations (so called free Ca for charge compensation of the negatively charged DNA) were then randomly placed ~6 Å from the surface using the Packmol package. ^45^ By placing the water molecules further from the surface this ensured the DNA was able to initially interact with the surface and not automatically displaced by the smaller and more mobile water molecules and cations. Perpendicular to the surface a large vacuum gap of ~60 Å separated the DNA/water from the unsolvated Ca surface. Across this gap a repulsive barrier was added to ensure water molecules were unable to desolvate from one calcite surface and move to the vacuum surface.

### DNA fragmentation in bulk experiments

#### DNA adsorption

200 uL elution buffer EB(Qiagen) was added to 20 mg of calcite powder (Sigma) and after sonication for 15 min. one ug of 1000 bp DNA fragment (Thermo Scientific™) was added in 300 uL EB buffer to the calcite samples (C1, C2, C3). The samples and two negative controls containing only calcite powder with no added DNA (samples 5 and 6) were incubated on a rotor for 4 hours. To mimic the AFM experiments we added we added 250 ul of molecular biology grade water (BioNordika) to disequilibrate the system allowing for slight DNA desorption and re-adsorption. After one hour of equilibration, we added another 250 ul of water and incubated them overnight. The samples were centrifuged for one minute at 13000 g, and the supernatant was extracted and used to measure the equilibrium concentration on a Qubit dsDNA High-Sensitivity assay (HS)(Thermo Fisher Scientific). None of the extracted supernatants had a measurable DNA. Finally, we washed the samples with one mL of EB buffer and centrifuged the samples for one minute at 13000 g. We retained the supernatant and measured the DNA concentration of the wash solution on Qubit. Only the wash solution from sample C2 had a measurable amount of DNA, which was 0.0134 ng/ul (13.4 ng total).

#### DNA extraction

After adsorption, samples were incubated overnight in one mL 0.5 M, pH 8 EDTA buffer (Invitrogen™) to release the DNA bound to calcite. In this step, we added positive control (samples 6, 7) where one ug 1000 bp DNA fragment was added to one mL EDTA and incubated overnight. All calcite-DNA samples (1-3) and negative controls (just calcite) (4-5) dissolved in EDTA during the incubation time. To purify the released DNA from EDTA, we desalted the samples using 10kDA Amicon® Ultra-4 Centrifugal Filter Unit (Millipore). We used a centrifuge with a swinging-bucket rotor at 4,000 × g for ~ 35 min until ~100 uL was left. Then we added three mL of molecular biology grade water and centrifuged the samples until ~100 uL was left. We repeated this step twice. Afterward, we measured the DNA concentration on Qubit dsDNA Broad-Range assay (HS)(Thermo Fisher Scientific) and send the samples for fragment analysis on Fragment Analyser (5300, Agilent) at the GeoGenetics sequencing core, University of Copenhagen.

## Results

### AFM imaging

We applied the same experimental procedure to both mica (negatively charged) and calcite (positively charged) surfaces to compare the DNA conformation and adsorption behaviour between the two types of surfaces. We applied 3 different 10 mM solutions of NaCl, MgCl_2_ and NiCl_2_ leading to two different ionic strengths: 10 mM, 30 mM and 30 mM and three distinct ionic potentials: 1, 12.5, and 16 respectively. When NaCl was used as the background electrolyte the DNA, as seen as thin white lines on the surface, did not adsorb to the mica surface (Fig. 1a). With MgCl_2_, the full plasmid adsorbed and generally in a ring form or with one or two coils (Fig. 1b). When NiCl_2_ was used, the DNA adsorbed as supercoils and the circular rings were not observed (Fig. 1c), indicating the stronger adsorption is preventing relaxation of the structure. Our results on DNA adsorption on mica agree with previously reported data (Zhai et al 2019).

**Figure 1.**
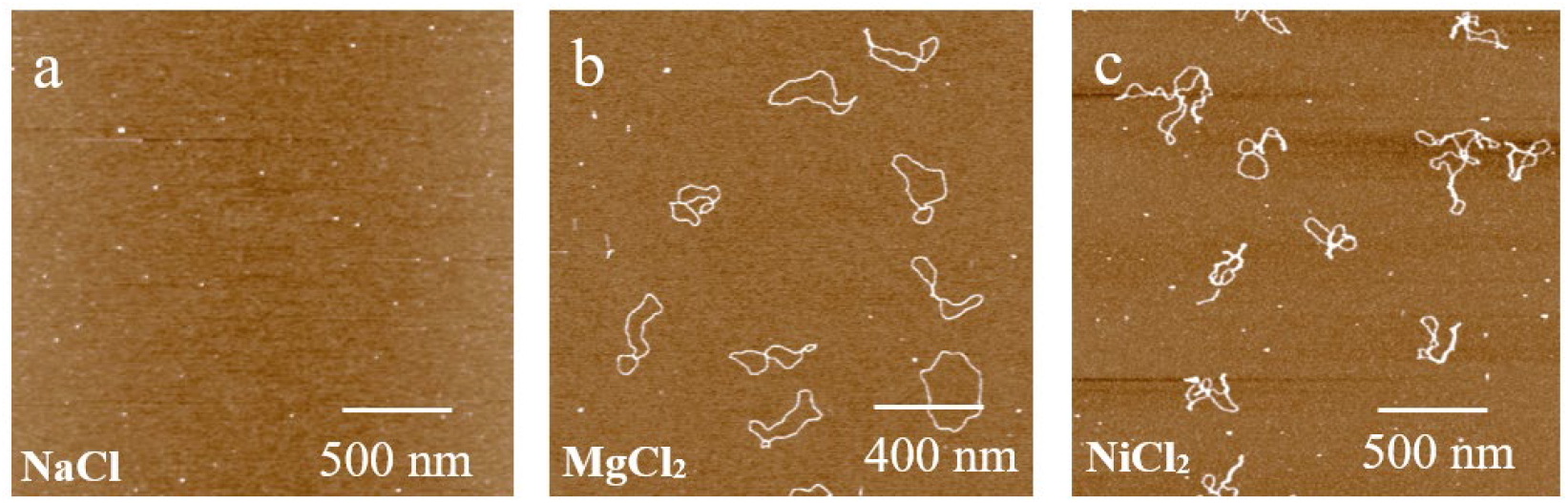
AFM images of mica showing the conformation of adsorbed plasmid DNA with different background ions: a) NaCl, b) MgCl_2_ and c) NiCl_2_.

Calcite is slightly water soluble and during the adsorption process, the DNA was added to calcite using a solution undersaturated with respect to calcite. This procedure causes a slight dissolution of the calcite surface resulting in the formation of etch pits (Fig 2a). The occurrence of edge sites surrounding the etch pits enabled us to assess if the DNA prefers such sites or accompanying terraces. The DNA adsorbs predominantly at the step edges with a clear preference for the acute step edges (Fig 2b). A part of the plasmid ring reaches across terraces to bridges to adjacent steps for maximising the plasmid’s interaction with step edges. The adsorption behaviour highlights that the interaction between DNA and the step edges is more favorable than between the terrace sites. A plasmid that showed preference toward adsorption on a terrace was not observed. No electrolytes were added to the experiment but in contrast to the solutions used in the mica study above (Fig. 1), there is always calcium present in the calcite system as calcite is a sparingly soluble salt and its surface will dissolve in calcite-undersaturated solutions. Although the resulting calcium concentration is much less than the 10 mM used for the background electrolyte, we cannot, from Fig. 2b rule out any effects that calcium ions in the solution have on DNA adsorption.

**Figure 2.**
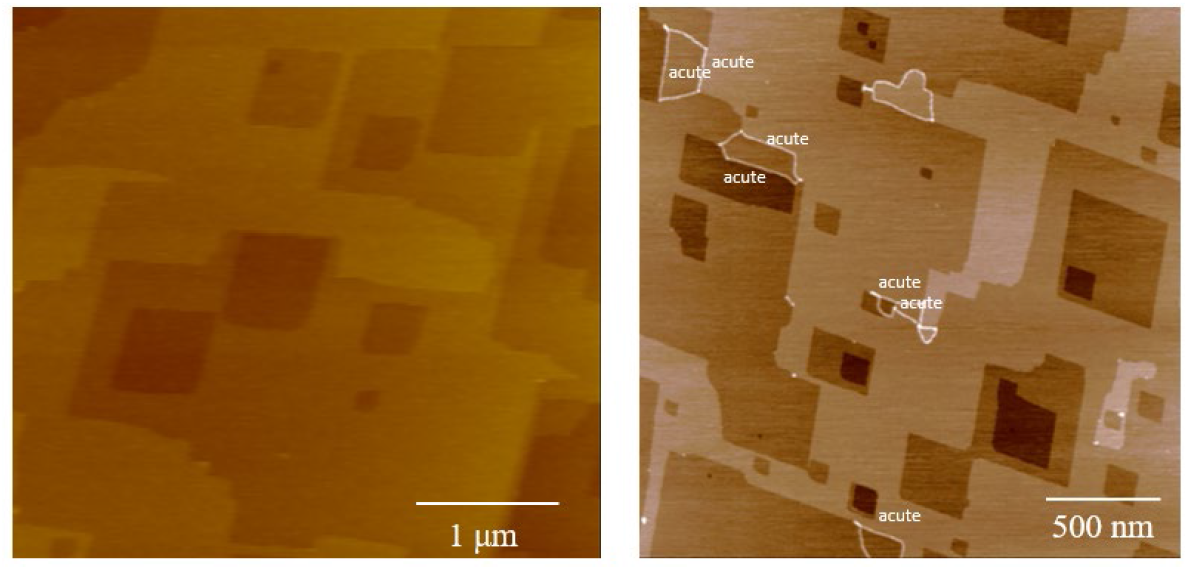
a) Freshly cleaved calcite after 2 min. exposure to the buffer solution used when adding the DNA to the surfaces. The image show step edges on steps and etch pits b) calcite surface displaying etch pits and adsorbed DNA. The etch pits morphology are determined by the underlaying orthorhombic structure of the calcite crystal which is oriented such that the obtuse corners is oriented south-west and the acute corner is oriented towards north-east. The DNA is visible as white (highestpoints in the image) lines lining some of the steps.

Applying the same range of background electrolytes as for mica (Fig. 3) we do see plasmid adsorption to calcite with the NaCl electrolyte (Fig.3a). We were not, however, able to discern any differences in DNA adsorption to calcite with either the added NaCl or MgCl_2_ background electrolytes (Fig. 3a,b). In the experiments where NiCl_2_ was used, we observed twisting of the helix (Fig. 3c-d) but the plasmid was still confined to the step edges and hence, probably did not have the freedom to coil as seen on the mica surface. Similar to Fig. 2b, we observed no adsorption where the majority of the DNA was bound to the terraces. This indicates that the steps, and hence the mineral surface charge density, is the determining factor controlling DNA adsorption to calcite and that the ionic potential of cations does not play as important a role as seen for DNA adsorption to mica.

**Figure 3.**
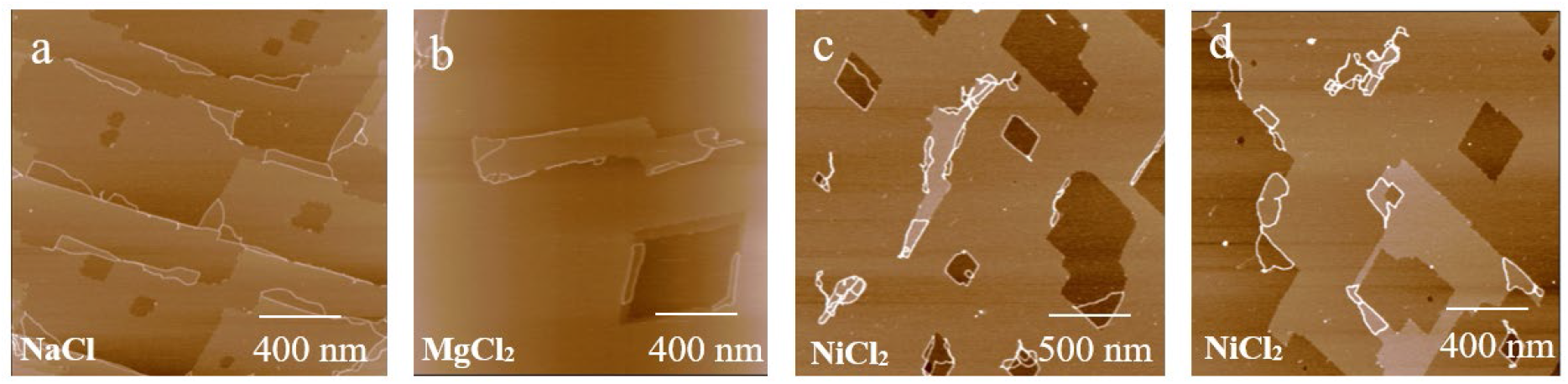
AFM images of calcite showing the conformation of adsorbed plasmid DNA with different background ions: a) NaCl, b) MgCl_2_ and c,d) NiCl_2_.

### MD Simulations

#### (10.4) terraces

Overall, the lack of DNA adsorption on the calcite terraces, as observed with AFM, are confirmed and further explained by the MD simulations. At the beginning of the simulations of DNA with the terrace (10.4 surface labelled configurations 1-6) the separation between the centre of mass of the DNA (i.e. the approximate centre of the cylindrical DNA) and the calcite surface is ~11 Å. Given the radius of the DNA cylinder is ~9 Å, this relates to parts of the DNA molecule being close or in direct contact with the surface. As the simulations progress the DNA moves perpendicularly away from the surface by an average of 1.96 Å as evidenced by the change in the centre of mass to surface separation (Fig. 4). The final distance between the surface and the center of the DNA molecule shows that the DNA moved above the tightly bound water layer on the calcite surface. The change in distance leaves a few direct interactions between the DNA and the calcite surface. Table 1 lists the direct interactions between the calcite surface and DNA molecule during the simulation and in only three of the six configurations the DNA is able to penetrate the water layer and make direct interactions with the (10.4) surface. In two such cases (configurations 2 and 5) oxygen atoms from the phosphate groups bind to calcium cations in the surface and in one case (configuration 3) part of the adenine base interacts with the carbonates of the surface. Examples of these interactions (for other cases) can be seen in Fig. 5 and the interactions are generally present during the whole of the simulation implying they are stable. Configurations 2 and 5 show the least perpendicular movement away from the surface.

**Figure 4:**
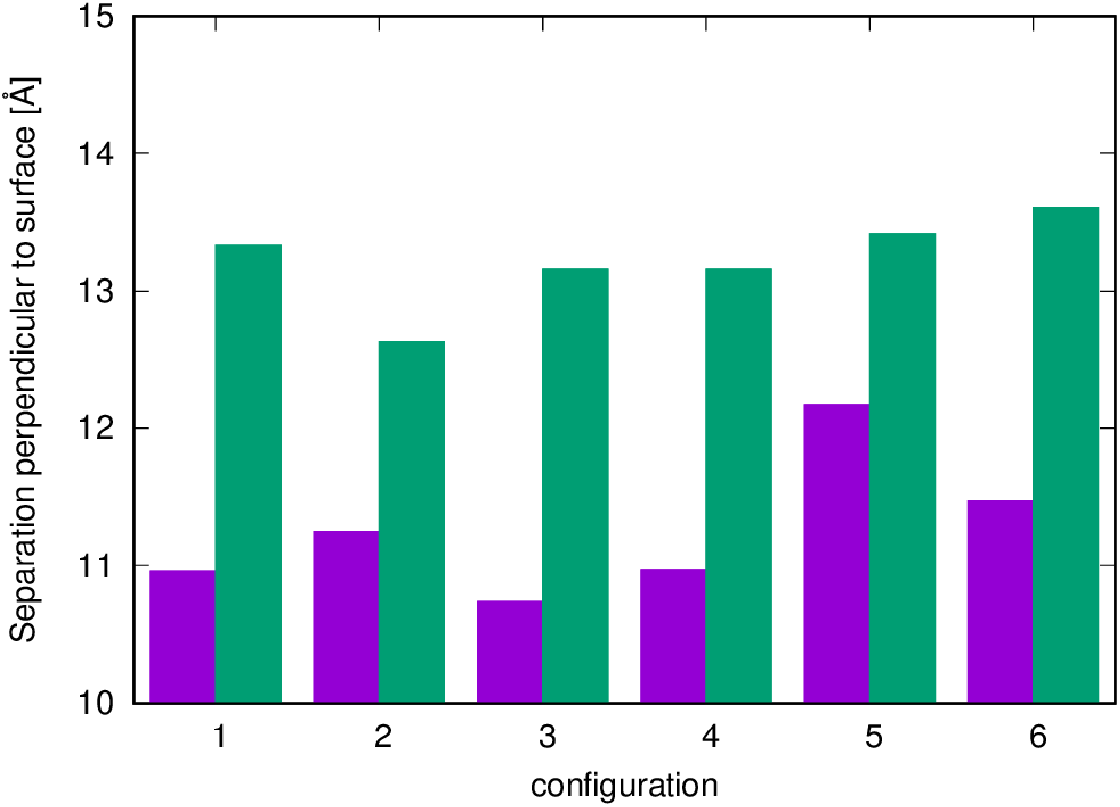
Separation perpendicular to the surface between the centre of mass of the DNA molecule and the calcite surface. Initial position (purple) vs final position (green). DNA has a radius of 9 Å.

**Table 1:** Change in H-bonding between residues in the two separate helices (from an average total of 21.8±1.95) and list of the number of interactions between surface calcium with O in DNA, DNA NH_2_-O(carbonate) in the surface and solvated calcium with O in DNA. Averages calculated over the last 500 ps of simulation. An interaction between DNA and surface or ions is defined as a maximum separation of 3 Å. An H-bond is defined as occurring between H and either an N or O atom at maximum separation of 2.1 Å.

| **Terrace (10.4)** | **H-bonds**<br>**change** | **Surface**<br>**Ca-O** | **NH<sub>2</sub>- O(carbonate)** | **Free Ca-O** |
| --- | --- | --- | --- | --- |
| 1 | -1.6 | 0.0 | 0.0 | 0.0 |
| 2 | +0.7 | 5.0 | 0.0 | 0.0 |
| 3 | -2.9 | 0.0 | 1.6 | 0.0 |
| 4 | -0.1 | 0.0 | 0.0 | 0.0 |
| 5 | +0.5 | 1.0 | 0.0 | 0.0 |
| 6 | -0.3 | 0.0 | 0.0 | 0.0 |
| **Acute carbonate +**<br>**obtuse calcium** |  |  |  |  |
| 7 | -3.1 | 3.0 | 0.0 | 0.0 |
| 8 | -1.9 | 0.0 | 0.0 | 0.0 |
| 9 | -5.9 | 1.8 | 0.0 | 0.0 |
| 10 | -3.5 | 0.0 | 0.0 | 0.99 |
| 11 | -4.8 | 0.0 | 0.0 | 0.91 |
| 12 | -0.7 | 0.0 | 0.0 | 0.0 |
| **Acute calcium +**<br>**obtuse carbonate** |  |  |  |  |
| 13 | -1.8 | 0.0 | 0.0 | 0.0 |
| 14 | -9.4 | 0.0 | 4.2 | 0.0 |
| 15 | -13.6 | 0.0 | 2.0 | 0.0 |
| 16 | -6.1 | 0.8 | 4.1 | 0.0 |
| 17 | -10.9 | 1.0 | 4.0 | 0.0 |
| 18 | -1.8 | 0.0 | 0.0 | 0.0 |

**Figure 5.:**
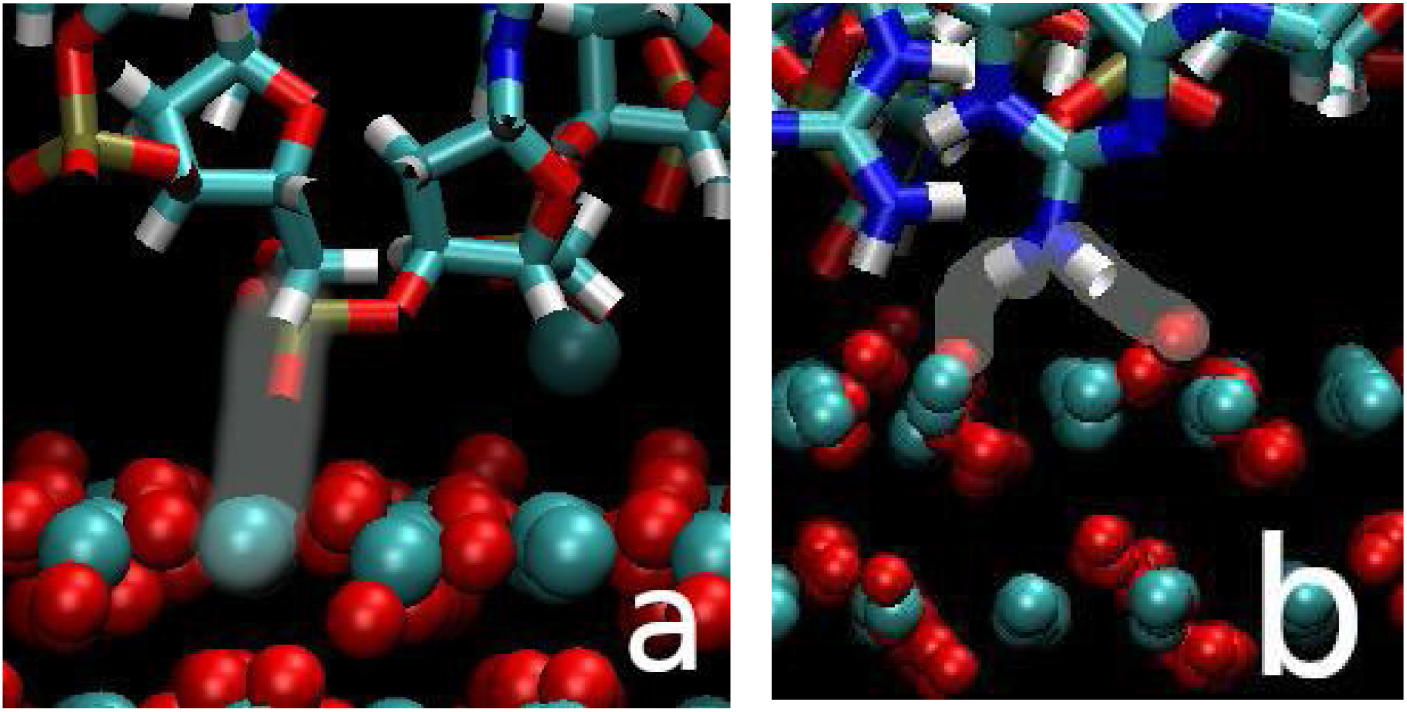
Example images of the direct interactions between the calcite surface and the DNA molecule. The interactions are highlighted to make them more visible. (a) Showing the O of the phosphate group and the Ca of the calcite as seen in configurations 2 and 5. (b) Showing the H of a NH_2_ group with O of the carbonate as seen in configuration 3. Key: O (red), H (white), C (cyan), Ca (cyan), N (blue), P (gold). Figure (a) taken from obtuse calcium step (configuration 7) and (b) taken from acute calcium step (configuration 17) acute step.

There are a total of 26 unique H-bonds normally present between the two strands in the double helix. Over the course of the simulation those H-bonds will break and reform due to structural fluctuations related to the normal thermal motion. When the DNA is run in a neutralised waterbox with no surface present an average of 21.8 H-bonds (with a standard deviation of 1.95) are present at any one time in the simulation. When placed in contact with the surface we generally see small changes in the number of H-bonds and the DNA structure is largely unaffected by the surface (see Table 1). The changes are small shifts in the lifetime of the H-bonds (full details found in the appendix) giving an overall reduction in the number of H-bonds.

The structure of the two chains still resembles the entwined double helix and the disruption is considered minor. Only in configuration 3 are more significant changes observed. In this configuration an adenine base comes out of the backbone and H-bonds to a carbonate group in the calcite surface. The loss of the base from the chain does not lead to a complete breakdown of the DNA structure but does significantly disrupt three of the base interactions (2 between cytosine number 10-guanine number 2 and 1 between thymine 11-adenine1) which leads to a net loss of ~3 H-bonds (Table 1, configuration 3). The net loss is within 2 sd of the mean, and hence could be described as a small disruption to the structure.

The charge-neutralising calcium that is free in the solution do not get involved in the binding process and none of them form interactions with the DNA (Table 1). Direct interactions between free calcium cations and oxygen atoms in the DNA molecule would break up the tightly bound solvation shells around the cations and are therefore unlikely to be energetically favourable. This suggests that the free ions are not key to the binding which agrees with our AFM experiments as the choice of electrolyte did not dictate whether binding occurred or not. The general infrequent interactions between the free Ca and the DNA agrees with our AFM where we observed little evidence of clumping or aggregation of the DNA molecules which is normally reported in liquid with divalent cations (Duguid and Bloomfield 1995).

Between the six different helix configurations at the flat terraces the average energies vary by only 75 kJ/mol and we do not find any correlation with the number of interactions with the surface. Configuration 3 displays the highest energy and is where the adenine base has twisted out of the helix suggesting that the energy gain associated with the surface binding does not outweigh the energy loss from the structural changes in the DNA molecule.

#### (10.4) Steps

The step structures used are shown in supplementary Fig. S2. As observed in the AFM images, the stepped surfaces show different adsorption behaviour than the terrace. The DNA strand started parallel to the edge of the periodic steps enabling us to measure the separation parallel to the surface from the centre of the DNA chain to the calcium and carbonate exposed step edges. At both surfaces we see a tendency for the DNA molecule to move towards the calcium exposed step (Fig. 6). Due to the geometry of the calcite, the calcium exposed step edges have a block of positive charge which attracts the negatively charged DNA backbone. The migration observed in the simulations demonstrates that the DNA will diffuse across the flat terraces and stop at the step edges. Examining Fig. 7 we can also notice that the DNA generally ends far closer to the acute calcium step edge than to the obtuse step edge. Both these observations are in agreement with our experimental results showing the DNA molecules adsorb at the step edges with a preference to the acute sites.

**Figure 6:**
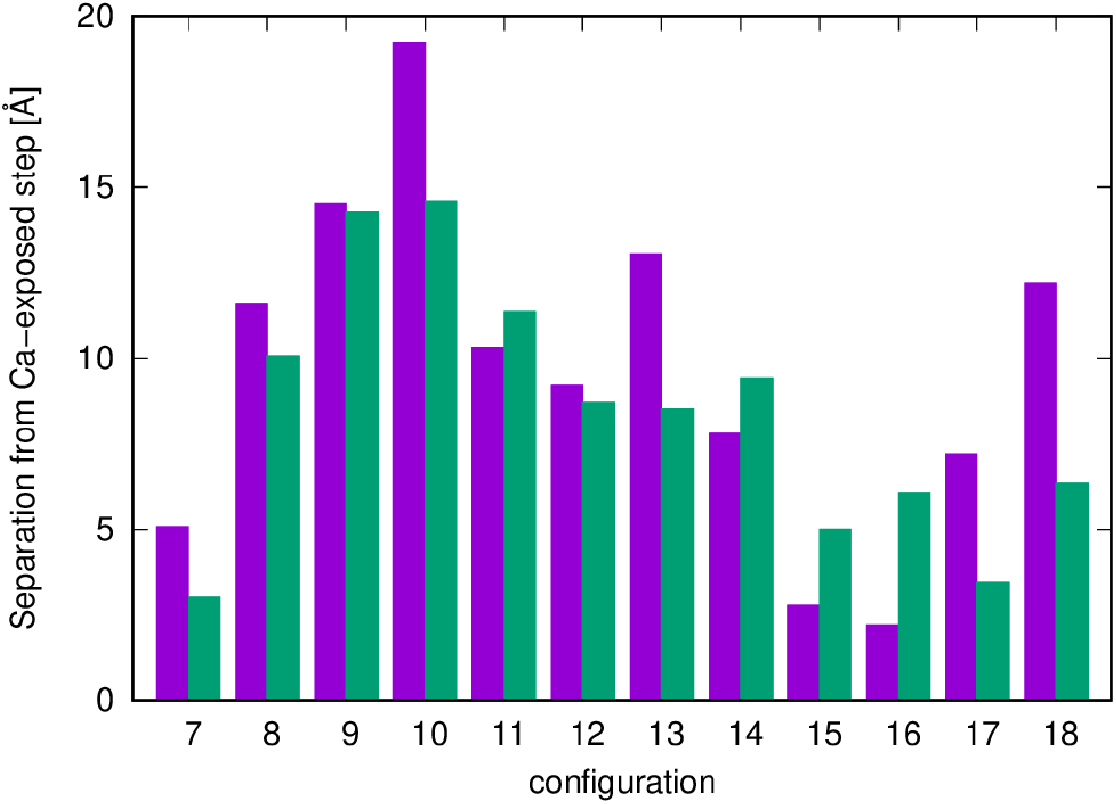
Separation parallel to the surface of the centre of the DNA molecule to the Ca-exposed step. Bars show starting separation (purple) and final separation (green). Configurations 7-12 are the obtuse calcium step surface. Configurations 13-18 are the acute calcium step.

**Figure 7.**
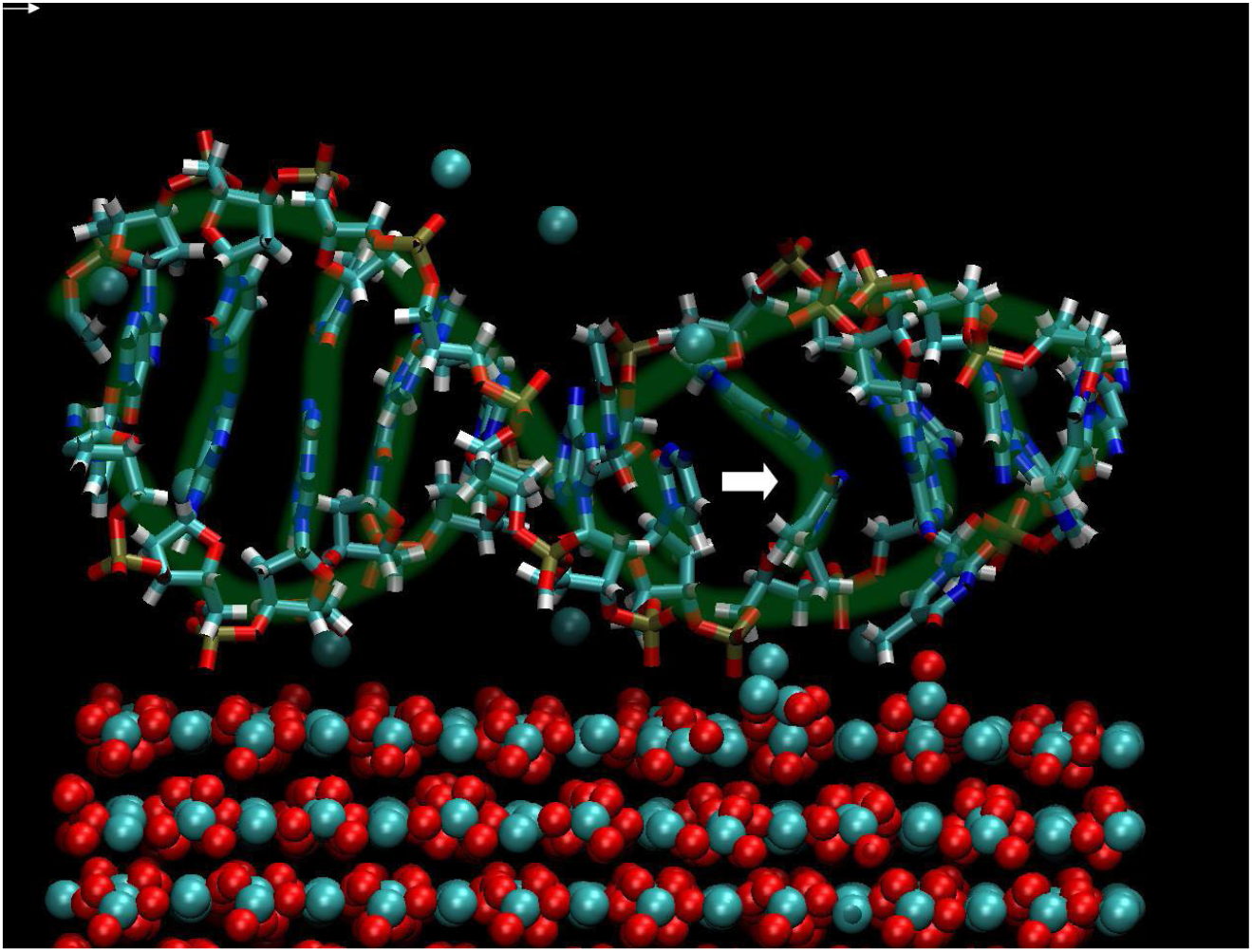
DNA on acute carbonate surface in configuration 7. The double helix structure has been highlighted in green to aid the eye. The arrow indicates a region where the double helix structure has lost some of its structure. Key: O (red), N (dark blue), Ca (cyan), P (gold), C (cyan), H (white), water not shown for clarity.

As also observed for the (10.4) terrace the bulk of the DNA strand resides above the organised water layers on the step surfaces. In half of the step based simulations there are no direct interactions between the DNA and the calcite surface. On the acute carbonate step surface, we observe two cases of the phosphate O backbone interacting with the Ca in the surface (configurations 7 and 9). On the acute calcium step we also see two cases of the backbone O phosphate with the surface (configurations 16 and 17). The remaining four direct interactions (configurations 14-17) come from the DNA bases that fold out of the helix on the acute calcium+obtuse carbonate step (Table 2). Of the bases that fold out of the DNA the same cytosine base is observed in all cases and the same adenine base is observed in three of the four cases suggesting that these are a particularly vulnerable part of the DNA sequence to folding out of the chain and binding. Two guanine bases are observed to interact directly with the surface in configuration 17. We never observe any thymine bases in contact with the surface which may be due to the random sequence and particular structure rather than their lack of NH_2_ amine groups. The NH_2_ group appears to be the major driver to the binding in the cases where the bases interact with the surfaces (Table 1) and remain relatively stable throughout the simulation.

**Table 2:** Listing the specific DNA bases that directly interact with the calcite surface for the acute calcium step system. Note bases are labelled with “a” for one helix and “b” for the other helix.

| Configuration | Bases |
| --- | --- |
| 14 | cytosine a1, adenine a3 |
| 15 | cytosine a1, adenine a3 |
| 16 | cytosine a1, adenine a3 |
| 17 | cytosine a1, guanine a9, guanine b2 |

The free calcium ions in the solution do not generally interact with the DNA (Table 1) but frequently cluster around the carbonate-terminated step due to the block of negative charge there (from the carbonate anions). In the simulation we observe two cases where the DNA does interact with the free Ca in solution (configurations 10 and 11). In these cases it is one loose Ca in the solution that interacts with the phosphate O. The interaction with the Ca ions does not operate as a bridge to the surface or similar feature.

Compared to binding at the (10.4) terrace the DNA undergoes a more significant structural change at the step edge. Even though only half of the configurations are binding to the surface directly, we see a significant loss of H-bonding in most of the configurations at the steps suggesting a general loss of inter-helix H-bonding rather than simple redistributions. This average loss of H-bonds is far larger at the steps (−0.62 on the (10.4) terrace, −3.32 at the acute carbonate and −7.27 at the acute calcium) and beyond a single standard deviation from the mean. Visual inspection shows a significant change in the DNA structure in some of the most affected configurations. From snapshots of the simulation progress we can identify particular cases of the breakup and obtain a better understanding of the process. Fig. 7 shows a snapshot of configuration 7 where several backbone phosphate O interact with the surface but there are no surface interactions from the bases. We have highlighted the double helix structure to show how the helix is largely retained during the simulation despite an average loss of 3.1 H-bonds which can be partially observed in the twisting of two bases within the helix (see the area highlighted with the arrow). Fig. 8a shows configuration 17 with a top down view (effectively looking at the plane of the calcite surface), it can be seen that one of the helix folds (the left and right hand ends of the image) has effectively opened and the bases moved apart (indicated by the arrows in Fig. 8a). The side on view in Fig. 8b shows how several residues move out of the core of the DNA molecule to bind to the acute calcium exposed step causing a major structural breakdown of the DNA (highlighted with the white arrows in Fig. 8b). We can also see that several bases are free but not interacting with the surface (left hand end of the image). Only in the central area of the DNA chain that can be seen more clearly in Fig. 8a is the helix preserved. In configuration 17 the DNA only has an average of 10.9 H-bonds left between its two helices, from the original 21.8 H-bonds. The results suggest that there is a strong attraction for the molecule at the step that is best achieved with the structural deterioration of the DNA molecule. The relatively low conformational freedom of the DNA molecule due to the periodic boundary conditions from the simulation likely prevents the full breakdown of the chain which may occur in a real system.

**Figure 8:**
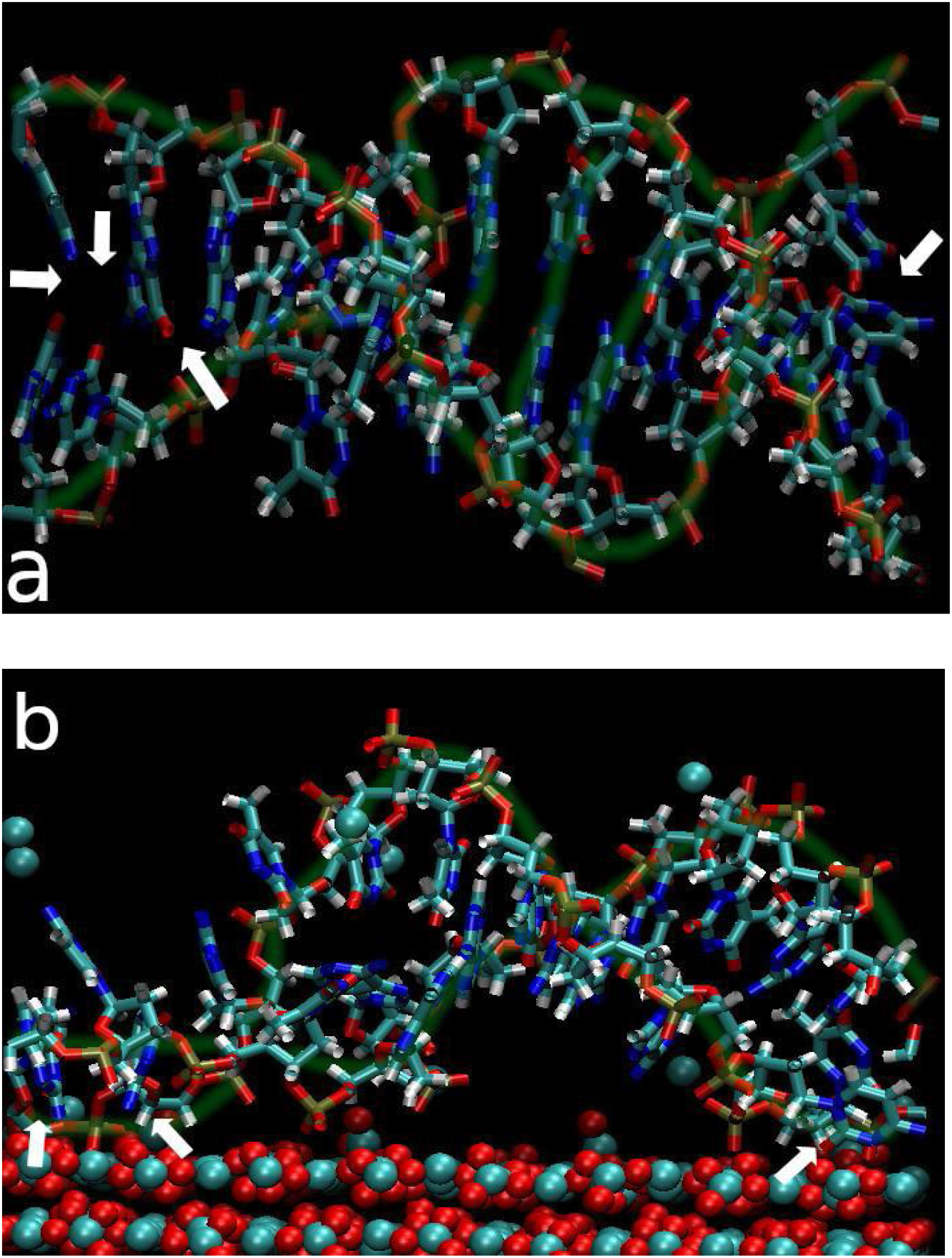
DNA on acute calcium step edge in configuration 17 showing the break out of the bases. The double helix structure has been highlighted in green to aid the eye. O (red), N (dark blue), Ca (cyan), P (gold), C (cyan), H (white), water not shown for clarity. (a) Top view looking down onto the surface (calcite surface not shown for clarity). (b) side view. Arrows are added to images to highlight breakdown of DNA structure in (a) and where bases bind to surface in (b).

The large positive charge at the calcium step edges will create a driving force to pull the DNA across the terraces towards the steps. It is likely that this positive charge build-up may cause deformations to the negatively charged phosphate backbone. The loss of H-bonds is larger at the step edges than on the terraces even when we see no evidence of the bases moving out of the helix (e.g. configuration 9 at the step). Once a base pair is separated from its opposite base pair then it is free to twist out of the chain and interact with the surface where it can make favourable interactions with both the step surface and terrace. The interactions between the DNA and the surface are most significant at the acute-calcium step edge. The acute step edge has previously been shown to have a more broken water structure compared to the obtuse step ^37,38^ meaning that binding of molecules is easier since there is less tightly bound water to displace. Binding of negative functional groups has also been highlighted in other molecular dynamics simulations.

In most cases the DNA was observed to move towards and interact with the calcium exposed step edges while avoiding the negatively charged carbonate exposed step edges. At the end of the simulation some of the free calcium in the simulation is often found in the vicinity of the carbonate exposed step edge as would be expected due to electrostatic interactions between the two. We further explored if calcium ions associated with a carbonate step edge would facilitate DNA binding. We made a set of starting configurations with all the free calcium within approximately 6 Å of the acute carbonate exposed edge. In this situation we observed that the DNA began ~13 Å from the calcium exposed carbonate step and did not move any closer to the acute-calcium step and instead remained close to the carbonate edge with the exposed calcium ions. This highlights that the presence of solvated calcium around the edge can neutralise the negative charge around the step and create a positive field that attracts the DNA molecule and therefore both acute step edges may favourably bind the DNA. It is difficult to isolate this interaction in the AFM experiments because we cannot know which step edges are terminated with carbonate versus calcium. There will be free calcium ions in the solution due to the dissolution of the calcite surface which could interact with the carbonate terminated step edge so this is a possibility in the real system. We should stress that we cannot observe ion structuring around the steps in the AFM experiments and we only explored one pH value in the experiments which will affect the step structure and ion concentrations. In the AFM experiments, however, we see DNA binding to the majority of the steps.

Comparing the energetics of DNA binding across the systems is not trivial due to the changes in DNA conformation during the simulation and the free ions in solution.. Fig. 9 shows the energy of each configuration with respect to the lowest energy configuration for that surface type (e.g. only comparing acute calcium and obtuse carbonate to other configurations with the acute calcium and obtuse carbonate) against the number of H-bonds present between the DNA helices. We can see a general trend is that as H-bonds are lost from the DNA then the energy of the total system is increased. This implies that those systems with more direct DNA-surface interactions will be higher in energy as the DNA loses its structural integrity. The significant changes in the DNA structure between the configurations also means that it is not possible to do a direct comparison of the adsorption energy between different surfaces as the DNA in the reference system is in a very different configuration (i.e. a perfect DNA molecule).

**Figure 9.**
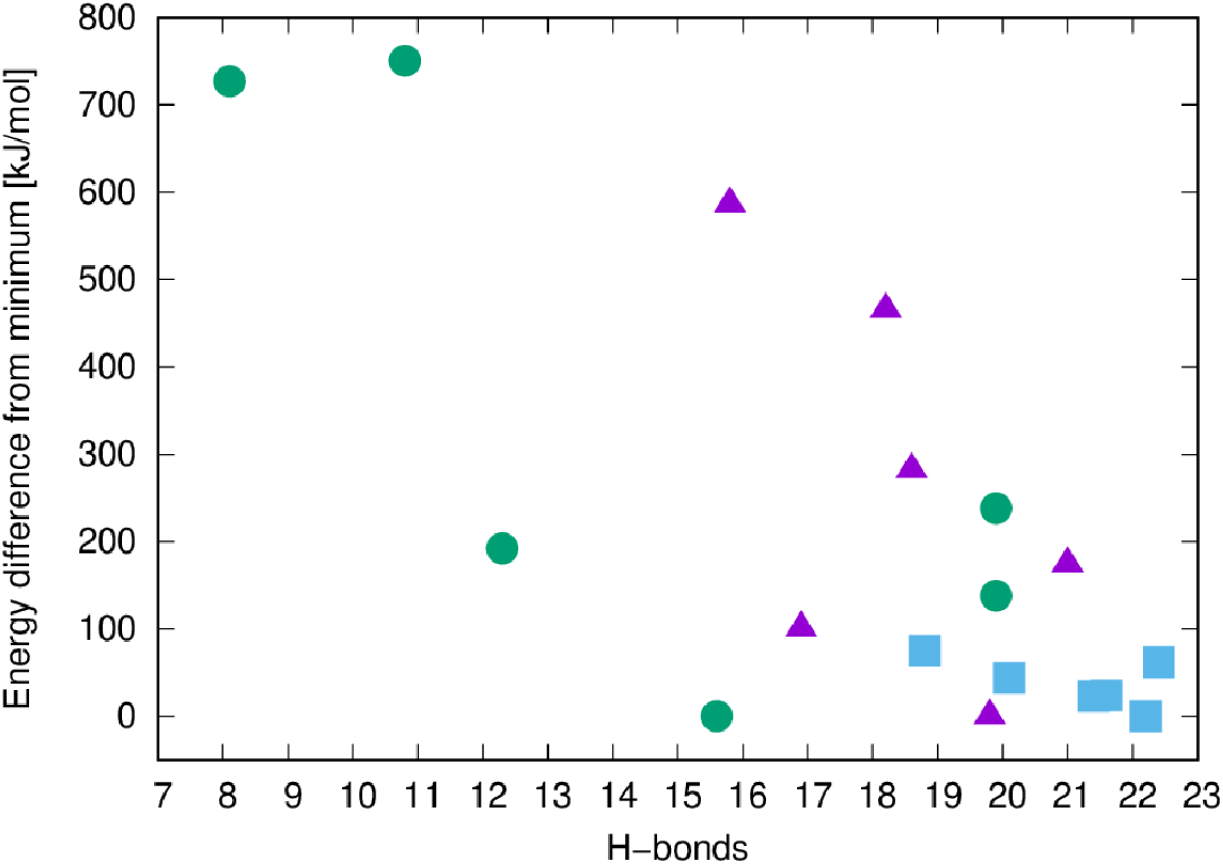
Relationship between number of H-bonds between residues in the two helices and the energy of the configuration with respect to the lowest energy configuration for that particular surface. Terrace (blue squares), Acute carbonate+obtuse calcium (purple triangles), Acute calcium+obtuse carbonate (green circles).

### Implications for sedimentary ancient DNA

When considered together, our experimental and modelling results highlight two key points that will influence the ability of extracellular DNA to be retained and preserved at a calcite or silicate deposit.

I. Retention of extracellular DNA at mineral surfaces is influenced by mineral charge density, solution composition and surface topography.
II. The potential fragmentation of bound DNA can be caused by strong interactions between the DNA and the mineral surface

We now discuss and explore these points with reference to DNA retention in sediments and access the preservation potential these two minerals can offer.

### Interplay between surface charge, solution chemistry and polymer charge on DNA conformation and adsorption

Surface charge density plays an important role in the behaviour of electrolytes and polymers near surfaces ^46,47^. As highlighted in Fig. 1, the ionic potential of the background ion plays a large role for how much of the DNA molecules was adsorbed. In this case the Na^+^ represented a cation with low ionic potential and Ni^2+^ a cation with high ionic potential. We fixed a positively, densely charged polymer (poly-L-Lysine) on a flat mica substrate to test if the high surface charge density would lead to supercoiling (Fig. 10). According to our predictions, the DNA supercoiled without the presence of charge dense background cations.

**Figure 10.**
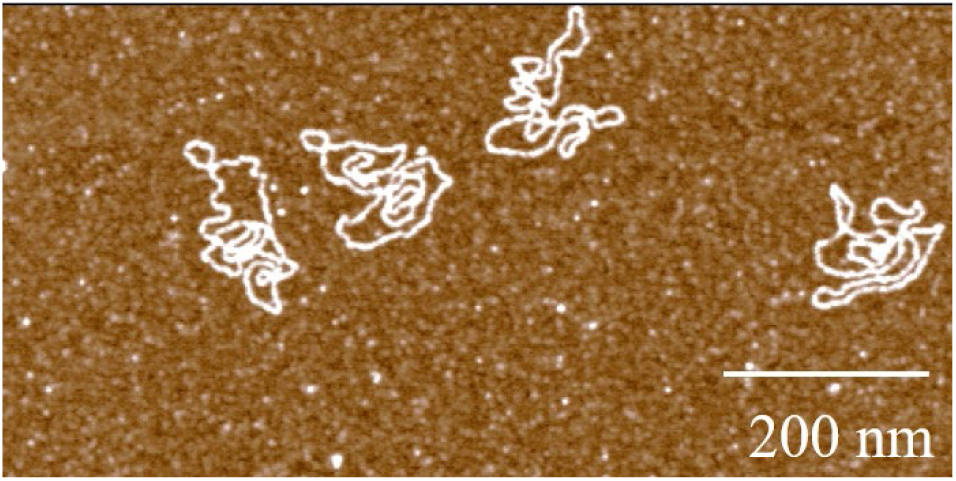
Adsorbed plasmid DNA on a Poly-L-lysine substrate. The DNA is supercoiled and was deposited with 10 mM NaCl as background electrolyte.

To probe the dependence between overall surface charge, ionic potential and solution concentration we used chemical force microscopy to measure the adhesion forces between a phosphate terminated AFM tip and calcite and mica surfaces (Fig. 11). The adhesion forces between the phosphate group and the calcite surface were, in general, higher than the forces between the phosphate tip and the mica surface.

**Figure 11.**
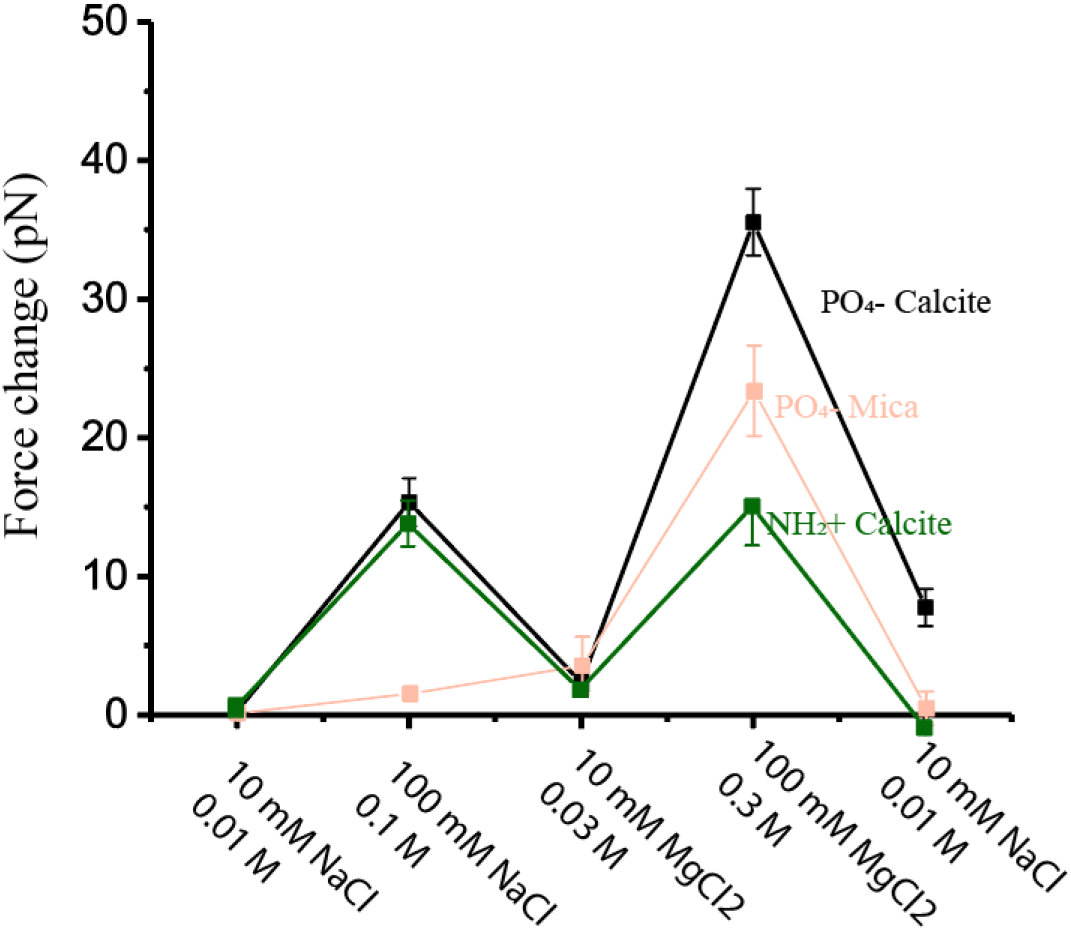
Adhesion force between a phosphate and an amine terminated AFMprobe and a calcite and a mica surface in a sequence of different concentrations of NaCl and MgCl_2_ solutions. The corresponding ionic strength is stated under the concentrations of each solution.

Across the applied sequence of solutions: 10 mM NaCl, 100 mM NaCl 10 mM MgCl_2_, 100 mM MgCl_2_ and 10 mM NaCl both minerals had the strongest interaction with 100 mM MgCl_2_, i.e. with a high concentration and a high ionic strength. The phosphate-mineral adhesion strength, however, showed different relative dependencies between the specific solutions. For mica the adhesion forces were only large when adding 10 mM Mg and far larger than the 100 mM NaCl indicating that at lower ionic strengths, charge density at the surface is more important than high salinity and the screening of the surface charge. For calcite the adhesion strength was highest in the two high concentration solutions indicating the thickness of the electrical double layer having a dominating role in DNA binding. It is worth noting that the calcite surface carries a “memory” of the past solutions, which causes an increased adhesion force for the 10 mM NaCl when measured after the 100 mM MgCl_2_ solution. Essentially the surface has not exchanged all the Mg ions with Na ions (the mica surface was re-cleaved prior to each force measurement and did not carry pre-adsorbed Mg ions). On the calcite surface the MD simulations indicate that the NH2 functional group of the base pairs is able to bind directly to the surface and therefore adhesion should be partially independent of the ion species in solution. The simulations also indicate that much of the DNA can struggle to bind directly to the surface through the water layers that would be disrupted by the presence of solution ions (Chen et al 2015) which could play a role for binding of the bases and associated structural breakdown. It is worth noting that the adhesion forces measured for calcite represent an average over both the steps and terraces. In general, the force distribution for calcite was broader than for mica which is expected as the calcite surface contains binding sites (steps and terraces) with different affinities for binding.

### Fragmentation

The MD simulations show that the binding on the acute-calcium (and potentially the carbonate) step could cause disruption of the helix because of the attraction between the base pairs and the surface through a NH-O(carbonate) interaction. We did not see evidence of the fragmentation in the AFM experiments despite a few smaller fragments observed among the intact plasmids. The small fragments could have been present in the stock solution and their presence is not evidence of fragmentation via binding. The AFM experiments were made in air and hence any fragmentation caused by the binding process would not be visible because the helix would still be bound to the step. To explore this further we used liquid cell AFM where we could record changes in DNA length and binding as a function of desorption and re-adsorption. After the initial adsorption of the DNA we slightly disrupted the equilibrium to induce dissolution of the calcite steps to initiate a desorption and readsorption process. The time it takes to set up scanning in liquid means that we cannot observe the initial desorption process i.e. when we reached the surface with the AFM tip initial restructuring had already occurred and the DNA was already fragmented. The fragmentation is evident in Fig. 12 which shows a time-resolved sequence of the same area displaying ongoing dissolution of the surface and DNA desorption and re-adsorption. The arrow in Fig. 12a) points to a DNA-free section of a step edge. In 12b) the space is filled, the terrace geometry has changed and there is a DNA-free side of the step (arrow). In 12c) that space is beginning to fill and the next step towards the left in the image (arrow) has a small amount of adsorbed DNA fragments. In 12d) there seems to be a movement of the DNA to that step (arrow) meanwhile the terrace is getting smaller. In 12e) only a small amount of DNA is left on the disappearing terrace edge and an increasing amount of DNA is observed on the step edge to the left (arrows). In 12f) the disappearing terrace is further dissolved and more than half of that step edge to the left is occupied by DNA. There is an empty section on the step edge (arrow) that has not yet been filled with DNA. In 12g) the empty section is filled up by a DNA fragment and only the upper part of the step edge is empty. In 12h) the step edge is filling up and there is a small section of the edge still free of DNA. In 12i) it is evident that the step is now dissolving and DNA is building up on the dissolving terrace edge (arrow). We cannot resolve if the DNA is getting increasingly fragmented because the fragments line up on the steps exploiting available space for adsorption. We observed no intact plasmids, even exploring the surface after we disrupted the equilibrium which supports that the binding does indeed facilitate fragmentation of the DNA.

**Figure 12.**
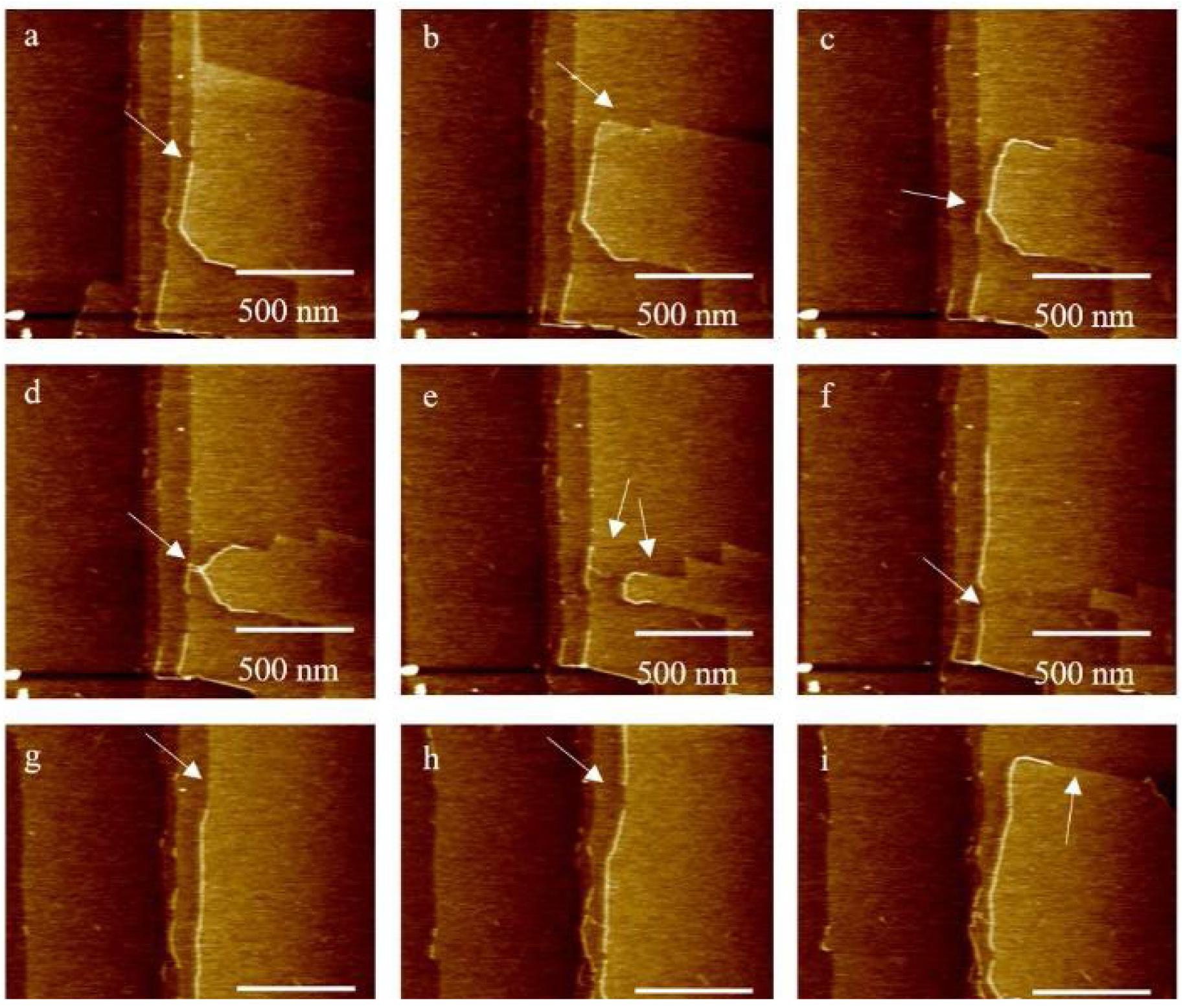
Time sequence of desorption and re-adsorption of DNA fragments as the calcite surface dissolves. There is 10-30 sec. between each frame.

We confirmed DNA fragmentation as a result of calcite binding via bulk DNA extraction experiments. In the AFM experiments we had a high surface to water volume (~1 cm^2^ surface and 40 ul liquid and 1 ng plasmid DNA with a further disequilibrium step by introducing 10 uL buffer). To quantify DNA adsorption, handle the extraction and ensure mixing we had a significantly lower surface to volume ratio in the bulk experiments. We added a double stranded 1000 bp DNA fragment to calcite powder and fragment analyser smear analysis (35 bp to 750 bp and 750 to 1000 bp) of the extract showed a DNA extraction efficiency for calcite samples was ~93, 90, and 29 %, respectively (Table S3). The extraction efficiency for positive control was ~78 and 98 %. Of the total amount of DNA extracted, short fragments (<750bp) accounted for 13, 16, and 10 % for calcite samples (sample 1-3) and 0.2 and 0.8 %, respectively, for the positive control (sample 6-7). These results suggest that most of the fragmentation happened on calcite, while only a small amount of fragmentation was caused by the extraction process. In the AFM experiment all the DNA images showed fragmentation. We ascribe the difference in fragmentation efficiency between the AFM and bulk experiments to the different solid to liquid ratio in the two setups. It should also be noted that AFM imaging does not provide a quantitative overview of the DNA as we did not attempt to quantify the DNA adsorption and nor is it possible to image the surface in its entirety.

### Preservation

Our results highlight that the adsorption process is an interplay between the adsorbed water layer, the charge density of the surface, surface topography and the polymer charge. For the negatively charged mica surface the ionic potential of the background ions plays a large role for the adsorbed DNA conformation and adhesion strength. For calcite the local charge density plays a role for the structuring of the associated water layer and the charge dense edge sites become the dominant sites for adsorption regardless of background ions. This behaviour is characteristic for outer sphere and inner sphere adsorption respectively. If, as in the case for mica, the DNA binds to the mineral through a cation bridge, the binding strength can be reduced by decreasing the concentration or the ionic potential of the solution. Such a change in solution conditions has been argued to essentially lead to desorption ^27,48^, and is unlikely to lead to fragmentation. In contrast, direct interaction between the DNA and the surface of calcite can create a strong association which becomes stronger with a decreased double layer. In the case of the acute calcium step, the interaction is strong enough to break up the chain and cause fragmentation.

If we translate our experimental and simulation observations to the retention of extracellular DNA in sediments, DNA adsorption to mica would be highly dependent on solution composition whereas calcite would be highly likely to tightly bind DNA in any environment. The strong association to calcite suggests that calcite adsorbed DNA is less likely to desorb if the composition of the background electrolyte is changed. In an open system such as soil or sediment, however, a change in solution composition would likely be followed by a shift in the calcite saturation of the solution, which could lead to desorption and re-adsorption with severe consequences for DNA preservation. The binding to calcite step edges is here shown to induce fragmentation and successions of de- and re-adsorption events would accelerate the fragmentation. In a cave environment, where the calcite equilibrium can be maintained within a speleothem (the water migrating there would be equilibrated by the time it reaches the mineral formation), any free DNA from a solution would have a higher chance of binding to mineral to the mineral surface. And even though the initial binding of DNA to calcite conjures some strand fragmentation, the long-term stabilisation of the strand in a relatively closed system such as some stalagmites would increase the potential for DNA preservation across timescales.

## Conclusions

Our results highlight that local mineral surface charge affects DNA adsorption. The interplay between DNA and background electrolyte becomes increasingly important for adsorption as the mineral surface charge decreases, such as for mica, suggesting ion bridging as an important adsorption mechanism. Additionally, a high positive surface charge density (calcite edges) facilitates the interaction with the DNA molecule but also causes a fragmentation of the double strand. Overall, the adsorbed conformation of the DNA is affected by both the adsorption process and ionic potential of the background ions, in particular if the molecule adsorption was not confined to topographical or charge dense sites but instead more freely to move on the surface.

Set in an environmental and depositional perspective, our data show that, a) for negatively charged minerals such as mica: the environmental setting is vital for both the adsorption process and also for understanding (post)depositional leakage and an important factor to take into account for determining DNA preservation in sediments and b) for positively charged minerals such as calcite DNA will be easily adsorbed and stored across time or through a range of environmental conditions provided the calcite is in equilibrium. The more fluctuations in calcite saturation the more fragmentation is expected. Consequently, fragmentation patterns are not necessarily a simple measure of age, but also, a function of events that disturb mineral solubility. The data presented here highlight that sediment mineralogy and environmental conditions during and post deposition can serve as a window for assessing DNA preservation potential at specific localities. Our study raises an important question: Is the eDNA we find and extract controlled by mineralogy and the environmental history of the deposit itself and do we know enough about these inter-dependencies and their role for DNA preservation to make interpretations on e.g. past biodiversity?

## Author contributions

CLF led and conducted the MD simulations and the MD interpretation and helped write the original manuscript. LD. conducted the AFM imaging, developed the experimental procedures associated and took part in the interpretations. OBAA made the CFM experiments, MZ led and conducted the extraction work. JQS contributed to the discussion and made one AFM image. MC provided context and helped draft the original manuscript. KKS conceptualized the study and designed the experimental work, interpreted the data and wrote the original manuscript.

## Acknowledgements

This work was supported by research grants from VILLUM FONDEN (00025352), the Danish Council for Independent Research (8123-00003A), and the Danish National Research Foundation (DNRF128). CLF acknowledges an EPSRC Programme Grant (grant EP/R018820/1) which funds the Crystallisation in the Real World consortium. OBAA acknowledges funding from the European Union’s Horizon 2020 research and innovation programme under the Marie Skłodowska-Curie grant agreement No. 892889. For the purpose of open access, MJC has applied a Creative Commons Attribution (CC BY) licence to any Author Accepted Manuscript version arising from this submission.

## Data availability statement

The data that support the findings of this study are available from the corresponding author upon reasonable request.

