## supplementary material for "Survival of environmental DNA in sediments: Mineralogic control on DNA taphonomy"

^3^ École nationale supérieure de chimie de Mulhouse, Université de Haute-Alsace,

3 Rue Alfred Werner, France^[1]^

**Supplementary Information**

Table S1 Ca-Oxygen (DNA) forcefield terms fitted using Schröder type method as described in Freeman et al. (2007).

| Atom 1 | Atom 2 | Type | A (kJ) | ρ (Å) | C (kJ Å^-6^) |
| --- | --- | --- | --- | --- | --- |
| Ca | OS^a^ | Buckingham | 155849.05 | 0.27151 | 0.0 |
| Ca | O^b^ | Buckingham | 220888.27 | 0.27151 | 0.0 |

^a^OS refers to ether type Oxygen which have two single bonds to chain forming atoms e.g. N, C or P.

^b^O refers to ketone type Oxygen which have a single double bond e.g. P=O, or N=O

| **Terrace (10.4)** | | |
| --- | --- | --- |
|  | DNA base H-bonds lost | DNA base H-bonds gained |
| 1 | - | - |
| 2 | - | - |
| 3 | Cytosine(10)-N:H-Guanine(2), Cytosine(10)-O:H-Guanine(2), Tyrosine(11)-O:H-Adenine(1) |  |
| 4 | - | - |
| 5 | - | - |
| 6 | - | - |
| **Acute carbonate+ obtuse calcium** | | |
| 7 | Tyrosine(7)-O:H-Tyrosine(4) | - |
| 8 | - | - |
| 9 | Cytosine(10)-N:H-Guanine(2), Cytosine(10)-O:H-Guanine(2), Tyrosine(11)-O:H-Adenine(1) | - |
| 10 | - | - |
| 11 | - | - |
| 12 | - | - |
| **Acute calcium and obtuse carbonate** | | |
| 13 | Tyrosine(7)-O:H-Tyrosine(4), Thymine(7)-H:N-Tyrosine(4) | - |
| 14 | Cytosine(1)-N:H-Guanine(11), Cytosine(1)-O:H-Guanine(11), Cytosine(1)-H:O-Guanine(11), Adenine(2)-N:H-Thymine(10), Adenine(2)-H:O-Thymine(10), Adenine(3)-N:H-Thymine(9), Adenine(3)-H:O-Thymine(9), Thymine(6)-H:N-Adenine(6), Adenine(8)-N:H-Thymine(4), Thymine(11)-O:H-Adenine(1), Thymine(11)-H:N-Adenine(1) | Thymine(6)-Adenine(6), Thymine(7)-O:H-Adenine(5) |
| 15 | Cytosine(1)-N:H-Guanine(11), Cytosine(1)-O:H-Guanine(11), Cytosine(1)-H:O-Guanine(11), Adenine(2)-N:H-Thymine(10), Adenine(2)-H:O-Thymine(10), Adenine(3)-N:H-Thymine(9), Adenine(3)-H:O-Thymine(9), Guanine(4)-H:N-Cytosine(8), Guanine(4)-H:O-Cytosine(8), Thymine(5)-O:H-Adenine(7), Thymine(6)-H:N-Adenine(6), Thymine(6)-O:H-Adenine(6), Adenine(8)-H:O-Thymine(4), Cytosine(10)-N:H-Guanine(2), Cytosine(10)-O:H-Guanine(2), Cytosine(10)-H:O-Guanine(2), Thymine(11)-O:H-Adenine(1) | Backbone-O:H-Guanine(11), Backbone-O:H-Guanine(11), Backbone-O:H-Guanine(11), Thymine(7)-O:H-Adenine(7), Guanine(4)-H:O-Thymine(9), Guanine(4)-H:O-Thymine(9) |
| 16 | Cytosine(1)-N:H-Guanine(11), Cytosine(1)-O:H-Guanine(11), Cytosine(1)-H:O-Guanine(11), Adenine(2)-N:H-Thymine(10), Adenine(2)-H:O-Thymine(10), Adenine(3)-N:H-Thymine(9), Adenine(3)-H:O-Thymine(9), Thymine(11)-O:H-Adenine(1), Thymine(11)-H:N-Adenine(1) | Backbone-O:H-Guanine(11), Backbone-O:H-Guanine(11) |
| 17 | Cytosine(1)-N:H-Guanine(11), Cytosine(1)-O:H-Guanine(11), Cytosine(1)-H:O-Guanine(11), Adenine(2)-H:O-Thymine(10), Thymine(6)-O:H-Adenine(6), Thymine(7)-O:H-Thymine(4), Thymine(7)-H:N-Thymine(4), Adenine(8)-H:O-Thymine(4), Adenine(8)-N:H-Thymine(4), Guanine(9)-O:H-Cytosine(3), Guanine(9)-H:N-Cytosine(3), Guanine(9)-H:O-Cytosine(3), Cytosine(10)-N:H-Guanine(2), Cytosine(10)-O:H-Guanine(2), Thymine(11)-H:N-Adenine(1) | Backbone-O:H-Guanine(11), Thymine(5)-O:H-Adenine(6), Guanine(9)-H:O-Backbone, Guanine(9)-H:O-Backbone |
| 18 | - | Backbone-O:H-Adenine(7), Thymine(7)-O:H-Adenine(6) |

Table S2: List of changes to H-bonding between the bases in the two DNA chains in the different surface binding configurations. See Figure SN for the structure of the DNA double helix. Each DNA base is labelled with a number to indicate its position in the chain and the H-bond is indicated with a “:”. In some cases H-bonds are made between the bases and the oxygen within the phosphate backbone of the chain. These are indicated with “backbone” label.

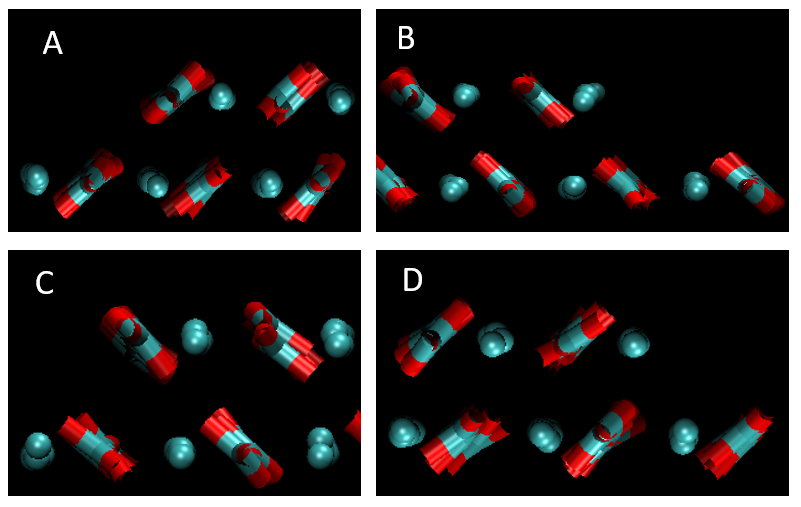

***Figure S2*** *Images of the different step edges and their terminations on the calcite surface. Note the angle of the carbonate molecule dictates the acute vs obtuse termination. (a) – obtuse carbonate, (b) obtuse Ca, (c) acute carbonate, (d) acute Ca. Key Ca and C (cyan), Oxygen (red).*

**Table S3:** Extraction data.

|  | Calcite and DNA | Calcite and DNA | Calcite and DNA | calcite | calcite | DNA | DNA |
| --- | --- | --- | --- | --- | --- | --- | --- |
|  | Sample | Sample | Sample | Negative control | Negative control | positive control | positive control |
| **Samples** | **1** | **2** | **3** | **4** | **5** | **6** | **7** |
| DNA conc. equilibrium concentration | - | - | - | - | - |  |  |
| DNA conc. Wash | - | 0.01 | - | - | - |  |  |
| DNA extracted volume uL | 105.00 | 100.00 | 110.00 | 85.00 | 105.00 | 100.00 | 130.00 |
| Smear analysis 35 bp to 750 bp (ng/uL) | 1.20 | 1.46 | 0.29 | 0.03 | 0.01 | 0.02 | 0.06 |
| Total amount DNA ng (35 bp to 750 bp) | 125.65 | 145.87 | 31.65 | 2.26 | 0.97 | 1.74 | 8.09 |
| % of 35 bp to 750 bp from total (35 to 1000 bp) | 13.44 | 16.17 | 10.60 | - | - | 0.22 | 0.82 |
| Smear analysis 750 bp to 1000 bp (ng/uL) | 7.71 | 7.56 | 2.43 | - | - | 7.81 | 7.55 |
| Total amount DNA 750 bp to 1000 bp (ng) | 809.13 | 756.42 | 266.95 | - | - | 780.68 | 980.94 |
| % of 750 bp to 1000 bp from total (35 to 1000 bp) | 86.56 | 83.83 | 89.40 | - | - | 99.78 | 99.18 |
| Total amount DNA ng ( 35 bp to 1000 bp) | 934.78 | 902.29 | 298.60 | 2.26 | 0.97 | 782.42 | 989.03 |
| Extraction efficiency % FA | 93.48 | 90.23 | 29.86 | - | - | 78.24 | 98.90 |

1. Current affiliation : L.C.I. S.à r.l, 2 ZAC Klengbousbierg, Bissen, Luxembourg
